# PTT-quant - a new method for direct identification and absolute quantification of premature transcription termination events, following the example of bacterial riboswitches

**DOI:** 10.1101/2021.06.18.448985

**Authors:** Piotr Machtel, Anna Wasilewska-Burczyk, Julian Zacharjasz, Kamilla Bąkowska-Żywicka

## Abstract

Regulation of gene expression by premature termination of transcription has been well described in all domains of life, including metazoans, yeast, plants and bacteria. Although methods for identification of such regulatory events by sequencing are available, the focused biochemical studies of the mechanism is hampered by lack of highly sensitive and accurate experimental methods. Here we propose a new method for absolute quantification of premature transcription termination events, PTT-quant. It is based on highly sensitive two-step digital droplet PCR protocol, coupled with normalized cDNA synthesis attained by site-specific pre-cleavage of investigated transcripts with RNase H. As a consequence, our method enables the reliable and sensitive quantification of both, prematurely terminated and full-length transcripts. By application of our method to investigation of transcriptional riboswitches in *Bacillus subtilis*, we were able to precisely measure the dynamics of SAM riboswitch induction, which turned to be ∼23% higher in comparison the results obtained without cDNA synthesis normalization.

## Introduction

The premature transcription termination (PTT) has been discovered a decades ago, but only recently it has been recognized as one of the major regulatory mechanisms leading to diversification of gene expression. Up to date, PTT has been reported in all kingdoms of life. In Eukaryotes, it can lead to synthesis of both, stable and unstable products. Depending on the gene region where termination occurs, two different types of PTT can be distinguished. The TSS-linked PTT is most likely related to low efficiency of full-length gene transcription by Pol II, resulting in high turnover within TSS regions [1,2]. This phenomenon lead to synthesis of substantial amounts of short, capped, mostly unstabel TSS-proximal RNAs of yet unknown function, which in a way similar to promoter upstream transcripts (PROMPT) contribute to exosome-sensitive RNAs. In contrast, the PTT occurring within the CDS region, is mostly related to occurrence of cryptic alternative polyadenylation (APA) sites. Among those, we can distinguish coding sequence polyadenylation (CDS-APA) and intronic polyadenylation (IPA) sites. Their employment is widespread, e.g. in mouse ∼40% of genes are subjected to PTT within the gene body [3]. The PTT events occurring closer to 5’ part of the gene tend to produce noncoding RNAs, whereas later PTTs result rather in synthesis of truncated proteins [4]. Most of such transcripts are capped and polyadenylated, playing important roles in cell metabolism. For example, PTT has been shown to form mRNA variants encoding soluble isoforms transmembrane T-cell co-stimulator CD46 [5].

In bacteria, PTT is mostly related to activity of RNA-based cis-regulatory mechanism. One of the best studied are riboswitches - highly structured mRNA domains with the unique capability of direct binding of small metabolites. Such binding event leads to changes in mRNA secondary structure and evokes regulatory effect [6]. Most riboswitches operate on the principle of negative feedback loops by their placement within the mRNAs encoding for the proteins which are involved in maintaining homeostasis of a recognized metabolite [7], including its biosynthesis [8], transport [9], or secondary metabolism [10]. Such regulation utilize a broad spectrum of mechanisms that act at various stages of gene expression, including transcription, translation, mRNA degradation, splicing or acting as sRNAs [11]. Importantly, the ligand:riboswitch interaction can silence [12] or activate gene expression [13]. Regulation occurs at the level of. To date, nearly 45 different classes of riboswitches were described [14]. Although riboswitches are considered as typical for bacteria [15-17], they have been identified among all domains of life [18-20].

Despite the wide wealth of regulatory mechanisms, the vast majority of currently known riboswitches act via PTT [21]. In the *apo* (unbound) riboswitch state, the expression platform adopts a conformation with a dominant anti-terminator structure, which allows for the undisturbed process of mRNA transcription and the production of full-length mRNAs. The binding of ligand (*holo* state) triggers the conformational alteration of a riboswitch, resulting in formation of a terminator hairpin structure, which in turn leads to PTT and the synthesis of short (truncated) nonfunctional mRNAs and gene silencing [22].

The identification of PTT events is in most cases fulfilled with employment of NGS-based techniques, being variations of 3’ directed RNA-seq. Quantification of the PTT efficiency using those methods is however affected by multiple experimental biases introduced during individual steps of cDNA library preparation, including differences in efficiency of 3’ adapter ligation, PCR amplification length bias, cDNA library composition, and others. Thus, identified PTT sites require further verification with employment of independent methods. Many of the currently employed approaches depend on quantification of the changes in full-length transcript levels, ignoring the concentration of truncated transcripts [23]. In this way the information concerning the actual activity of the PTT event is lost. Moreover, measured changes in transcript levels are the resultant of multiple cellular processes, including activity of RNA polymerase or RNA turnover [24,25]. Thus, to estimate efficiency of PTT event, both fractions of transcripts (full-length and truncated) should be quantified and their ratio should be estimated.

When aiming to accurate detect terminated and full-length transcripts, several issues must be taken into consideration. Routinely exploited hybridization-based techniques, like northern blot hybridization, although enable amplification-free estimation of mRNA levels, is hampered by several limitations, eg. the inability for controlling the efficiency of RNA transfer from the gel to the membrane, RNA-membrane binding, and differences in kinetics of detection probe hybridization to targeted nucleic acids [26] [27]. Moreover, northern blot analysis can cause RNA degradation during electrophoresis steps. All those issues low sensitivity and resolution, much lower in comparison to amplification-based techniques [28].

Thus, the most frequently employed technique for PTT investigation is RT-qPCR aimed at detection of both, short and full-length transcripts. Although this method provides high sensitivity, it is hampered by biases during cDNA synthesis. In one-step RT-qPCR protocol, same set of starters are used for reverse transcription and PCR reactions. Thus, the comparative analysis of prematurely terminated and full-length transcripts con not be performed from single RNA sample. The requirement of using two separate RNA samples, even if obtained from same RNA pool, could lead to measurement errors which influence the estimated PTT efficiency. Alternatively, two-step RT-qPCR protocol can be applied, where cDNA is synthetized using random hexamers as starters, and next, on single cDNA pool two PCR reactions can be performed. This approach however reveal a systematic bias towards overrepresentation of 5 ends of transcripts during cDNA synthesis [29]. The reason for this phenomenon is the fact that reverse transcriptase synthesize cDNA in the direction from the 3 to the 5 end (in relation to the transcripts); therefore there is a high probability that regions of mRNAs closer to the 3 ends will be converted into cDNA with lower frequency, compared to 5 ends (Figure 1). Thus, the results of quantification depend not only on amount of RNA, but also on the potions of the designed PCR primers within the transcript. The above effect introduce bias in relative quantification of prematurely terminated and full-length transcripts. It specifically concerns transcripts with significant length and could contribute to artificial overestimation of PTT transcripts.

**Figure 1.**
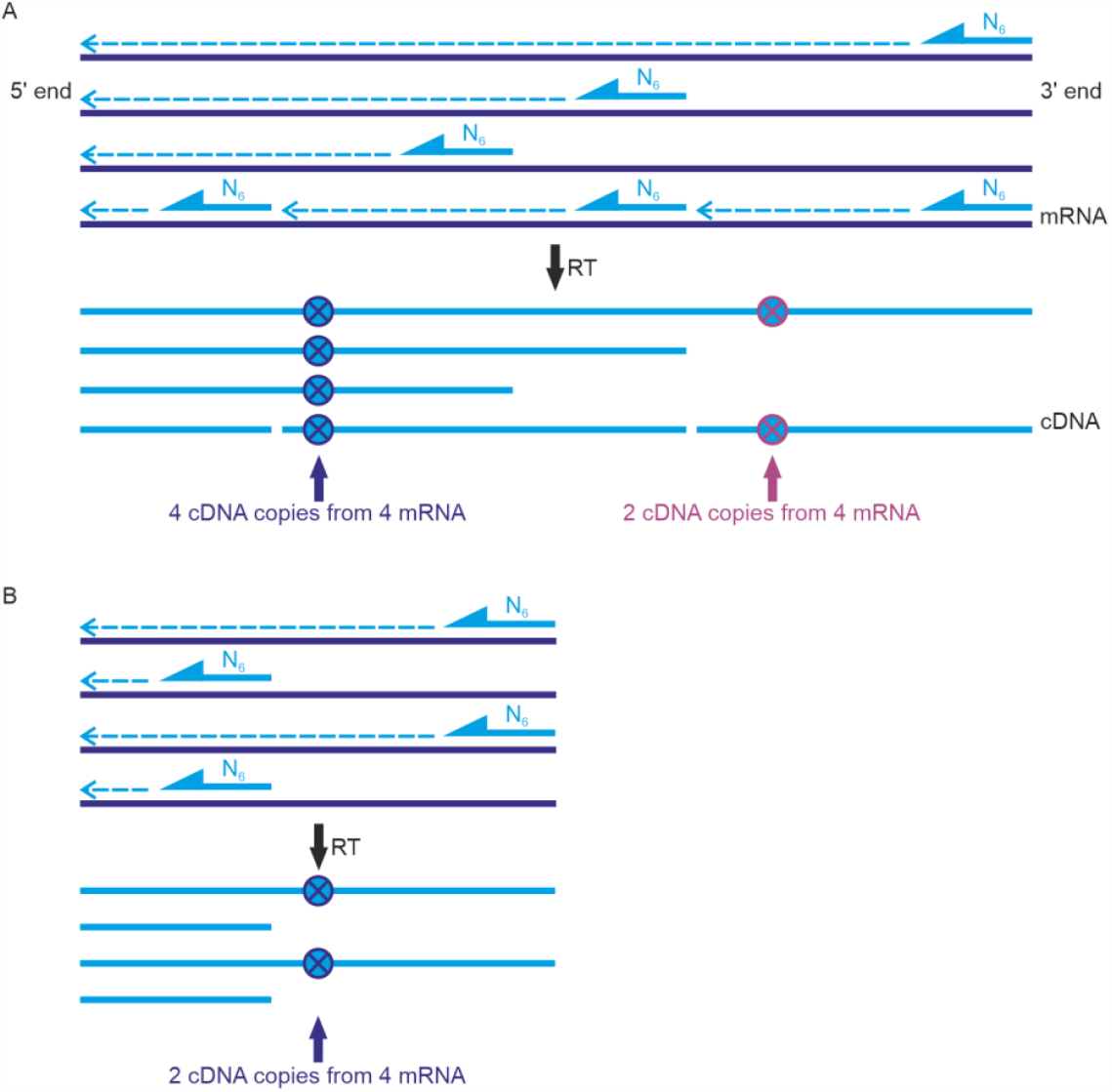
Dependence of the number of cDNA copies on distance from the 3 end of transcript. Full length transcripts (A) and terminated transcripts (B). Dashed arrows - reverse transcription reaction direction. N6 - random sequence hexamer.

Due to above limitations in currently available protocols, we decided to establish a novel method for quantification of premature transcription termination events, that will be flexible, accurate, sensitive and less prone to introduction of biases. As the model, we have employed SAM binding riboswitches in *Bacillus subtilis*. Bacterial transcriptional riboswitches are well suited for such task, due to relatively straightforward control over PTT activity controlled by ligand concertation. Also, the exact position the PTT event in the sequence can be easily estimated by identification of the attenuator hairpin following investigated riboswitch’s aptamer domain.

## Materials and methods

### Primers and DNA oligomers

Primers for the PCR (Table S1) were designed using Primer BLAST tool and *Bacillus subtilis* 168 strain genome sequence as a reference (NCBI database accession number AL009126.3).

### Bacteria strain and growth conditions

The *Bacillus subtilis* 168 strain was used in the study. For routine growth, the *B. subtilis* cells were grown on Sporulation Medium (SP) or ED minimal medium [30]. Bacteria were grown at 37°C to OD_600_=0.6. Methionine elimination was performed as follows: *B. subtilis* was grown to OD_600_=0.6, the pellet was centrifuged and resuspended in ED medium lacking methionine. The culture was continued for up to 4h.

### DNA isolation

Genomic DNA (gDNA) was isolated from 2 ml of *B. subtilis* bacterial culture with OD_600_from 0.8 to 1. The bacteria were centrifuged and a pellet was resuspended in 500 μl of cold DNA isolation solution (50 mM glucose, 2.5 mM Tris-HCl pH 8.0, 10mM EDTA pH 8.0, 2 mg/ml lysozyme,100U RNase A) and incubated for 10 min on ice. Then 50 µl of 10% SDS was added and the resulting mixture was incubated at 37°C for 10 min. gDNA was phenol/chlorophorm extracted. DNA quality and quantity were verified with NanoDrop spectrophotometer and gel electrophoresis.

### RNA isolation

RNA was isolated with GeneMATRIX Universal RNA purification kit (EURx), according to the manufacturer’s protocol. The RNA quality and quantity were measured with NanoDrop spectrophotometer and gel electrophoresis.

### PCR

PCR templates for *in vitro* transcription and *ex vivo* RNase H cleavage were amplified using DreamTaq PCR Master Mix 2x (Thermo Fisher Scientific). One microgram of *B. subtilis* genomic DNA was used as a template and the final concentration of primers equaled 200 pM each. The PCR products were purified with GeneJET PCR Purification Kit (Thermo Fisher Scientific) and their purity was verified on 1% agarose gel with SYBR Safe reagent (Thermo Fisher Scientific) and visualized on G-Box Chemi XR5 system (Syngene).

### *In vitro* transcription

*In vitro* transcription of PCR products was performed with MEGAscript™ T7 kit (Thermo Fisher Scientific) using a 45 nt-long primer (5’-TAATACGACTCACTATAGGGAGAATGAGTGAACAAAACACACCAC-3’). Wight microliters of PCR template was used, the final concentration of NTPs equaled 7.5 mM each. Samples were incubated for 4 h at 37°C. Subsequently, *in vitro* transcription products were purified with a GeneMATRIX Universal RNA purification kit (EURx) and visualized on a 10% PAA gel with SYBR Safe reagent (Thermo Fisher Scientific).

### RT-PCR

RT-PCR was conducted using SuperScript IV Reverse Transcriptase (Thermo Fisher Scientific). RNA molecules obtained *ex vivo* were used as templates. Synthesis of cDNA was carried out with the use of 2.5 μM random hexamer primers, 0.5 mM dNTP each and 1 ng of RNA template. The reaction mix was incubated at 65 °C for 5 min and on ice for 1 min. Subsequently, 4 µl of SuperScript IV RT buffer, 1 µl of 5mM DTT, 0.5 µl of RNaseOUT Recombinant RNase Inhibitor (40 U/µl) and SuperScript IV Reverse transcriptase (200 U/µl) was added. Samples were incubated in thermocycler at 23°C for 10min, 50–55°C for 10 min and at 80°C for 10 min.

### RNase H cleavage

DNA oligomers complementary to selected RNAs were designed using Primer BLAST tool (Table S2). 1 μg of transcript template was incubated with 100 pM to 5 µM DNA oligomers (depending on the experiment) for 20 min at 37°C. A hydrolysis of DNA-RNA duplex was performed using 1.25 U Ribonuclease H (Thermo Scientific).

### cDNA synthesis

For cDNA synthesis SuperScript IV Reverse Transcriptase transcription kit was used. The reaction was carried out according to the manufacturer’s protocol with the use of 50 µM random hexamers. To each experiment,100 ng of RNA was used as a template. The first part of reaction, containing RNA templay, random hexamers and NTPs was incubated at 65°C for 5 min and put on ice, followed by incubation at 23°C for 10 min, 55°C for 10 min and 80°C for 10 min with remaining reaction components..

### Droplet Digital PCR

ddPCR reaction was performed as previously described [31] with some modifications. ddPCR mix contained: 10 µl of QX200™ ddPCR™ EvaGreen Supermix (Bio-Rad), 4 µM of forward and reverse primers, 1 µL of cDNA in 20 µL total volume. The reaction mixture was emulsified into droplets using a QX100 Droplet Generator (Bio-Rad) according to the manufacturer’s protocol. PCR was performed according to the program: 5 min at 95°C; followed by 40 cycles of (30 sec at 95°C, 30 sec at 55°C, 45 sec at 72°C); then 2 min at 72°C, 5 min at 4°C, 5 min at 90°C and finally kept at 12°C. Samples were analyzed using the QX100 Droplet Reader (Bio-Rad) and analyzed with QuantaSoft software (Bio-Rad).

### RT-qPCR

RT-qPCR was performed as previously described [32] with some modifications. RT-qPCR mix contained: 4µl of 5x HOT FIREPol® EvaGreen® qPCR Mix Plus (Solis Biodyne), 200 nM of forward and reverse primer each, 5 µl of cDNA samples (dilution 1:50). Datasets were collected on an Applied Biosystems™ QuantStudio™ 6 Flex Real-Time PCR System (Thermo Fisher Scientific). The cycling conditions were as follows: 12 min at 95°C, followed by 40 cycles consisting of 15 sec at 95°C, 20 sec at 60°C and 20 sec at 72°C. Fluorescence signal data were collected during the 72°C phase of each cycle. The specificity of amplified targets was assessed by melting curve analysis from 55°C to 95°C (in 0.5°C increments, measuring fluorescence at each temperature) following the last cycle. The analysis showed the presence of only one specific product in each reaction. All primer pairs were tested with regard to amplification efficiency with 4x log10 serial dilution of a random cDNA sample in triplicates. All tested primers met the criteria of efficiency 90–110% and R^2^ > 0.985. The results were expressed as a relative quantity according to the equation RQ=2^ΔΔCt^.

## Results

### Protocol design

Among providing high sensitivity amplification-based techniques, reverse-transcriptase digital droplet PCR (RT-ddPCR) seem to be best suited for investigation of PTT activity, due to superior accuracy without need for standard curve estimation, obtained by absolute quantification of nucleic acids. It also enables the employment of the two-step protocol, consisting of initial reverse-transcription of total RNA with employment of random hexamers, followed by multiple target-specific ddPCR-based cDNA quantifications from single cDNA sample. This method is however biased by overrepresentation of 5’ RNA regions as a result of non-uniform cDNA synthesis, similarly to classical two-step RT-qPCR (see Figure 1). The severity of this bias observed in RT-ddPCR results has been shown by Abachin et. al [33]. During the optimization of RT-ddPCR workflow for detection of dengue virus, they investigated the efficiency of RT-ddPCR with employment of probes designed toward different regions of the viral RNA genome. The results demonstrated a clear drop in assay sensitivity for assays based on probes located closer to 3’ end, especially within 3’ UTR.

In order to eliminate the influence of the 3’ end distance on the number of reads (Figure 1), we decided to use endonucleolytic cleavage of the investigated full-length transcript with RNase H at the position defined by designed DNA oligomers, complementary to the sequences near the natural PTT site (Fig. **2**). As the result, we obtain two types of RNAs, first, representing 5’ part of the transcript and second, representing 3’ part, being extension beyond the investigated PTT site. Thus, quantification of the first one provide information about total expression of the gene, whereas second provide information about full-length transcription. By designing ddPCR probes for detection of both isoforms at the positions in the same distance from their 3’ ends, we are able to ensure unbiased efficiency of cDNA synthesis, for both probe sets, independently from measured transcript length.

**Fig. 2.**
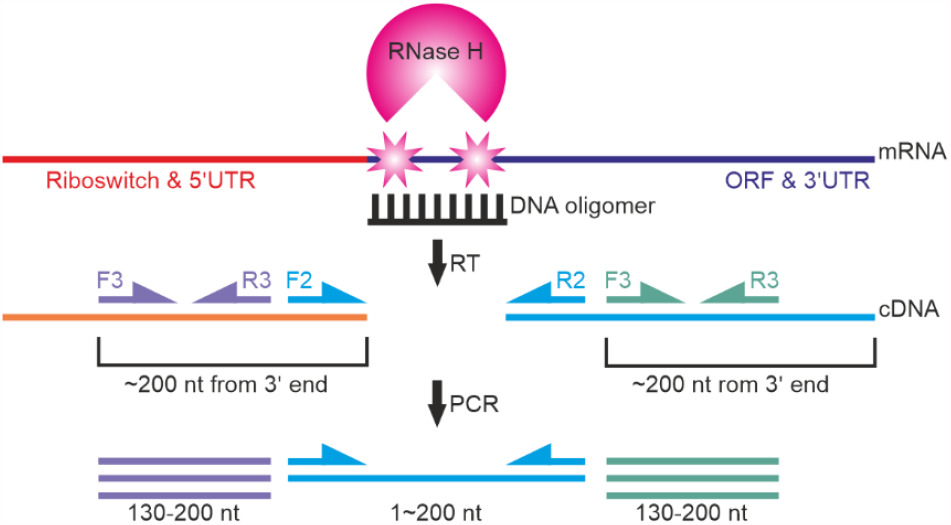
Schematic representation of a new method for the analysis of premature transcription termination transcriptional activity and primers/oligomer design. F – forward primer, R – reverse primer, FL – full-length products, T – total (PTT + full-length) products.

However, in our experiments, we have designed 3 pairs of primers for each investigated transcript. The first pair of primers (Fig. 2, primers F1 and R1) are complementary to the riboswitch aptamer sequence at 5 UTR, upstream of the PTT site. The second pair of primers (primers F2 and R2) flank the transcription termination site, whereas the third pair (primers F3 and R3) was designed to bind to ORF sequences. Primers 1 and primers 3 were designed to hybridize at the same distance from the 3 end of the RNAse H cleavage product and full length transcript, respectively. In this way, the reads from primers 1 can be used for quantification of to the sum of full-length and terminated transcripts, primers 3 - provide the information about amount of full-length transcripts, whereas primers 2 - provide control for the RNase H cleavage reaction. The difference between the number of the reads from the first and the third pair of primers gives the information on the number of truncated transcripts, which are produced as a result of the riboswitch-induced PTT activity.

### Optimization of efficiency and specificity of RNase H cleavage of *in vitro* transcripts

To optimize the efficiency and the specificity of RNase H cleavage we prepared a set of cleavage reactions *in vitro*. To directly observe the action of RNase H, in the first step we synthesized a model transcripts with the use of *in vitro* transcription reaction on PCR-amplified templates containing T7 promoter for RNA polymerase, coupled with a partial sequence of the *gyrA* gene composed of 5’UTR, and a fragment of an downstream ORF sequence. The RNase H cleavage site was defined by design of a complementary DNA oligomers, selectively complementary to the target RNA sequence near the site of native PTT caused by riboswitch activity (i.e. behind the terminator hairpin and poly-U tract).

Three sizes of DNA oligomers were tested (15 nt, 20 nt, 25 nt, Table S2) to determine the impact of the length of DNA oligomers on RNase H cleavage efficiency (Figure 3). The analysis showed no significant differences in the cleavage efficiency. In case of all three oligomers, the cleavage efficiency was constant. The only visible differences concerned the size of the analyzed products – as expected from RNAse H cleavage. The reason is the fact that RNase H induces more than one nucleolytic cleavage site, along the whole RNA:DNA heteroduplex. Therefore, oligonucleotides of ∼20 nt in length were used for further analysis. The

**Figure 3.**
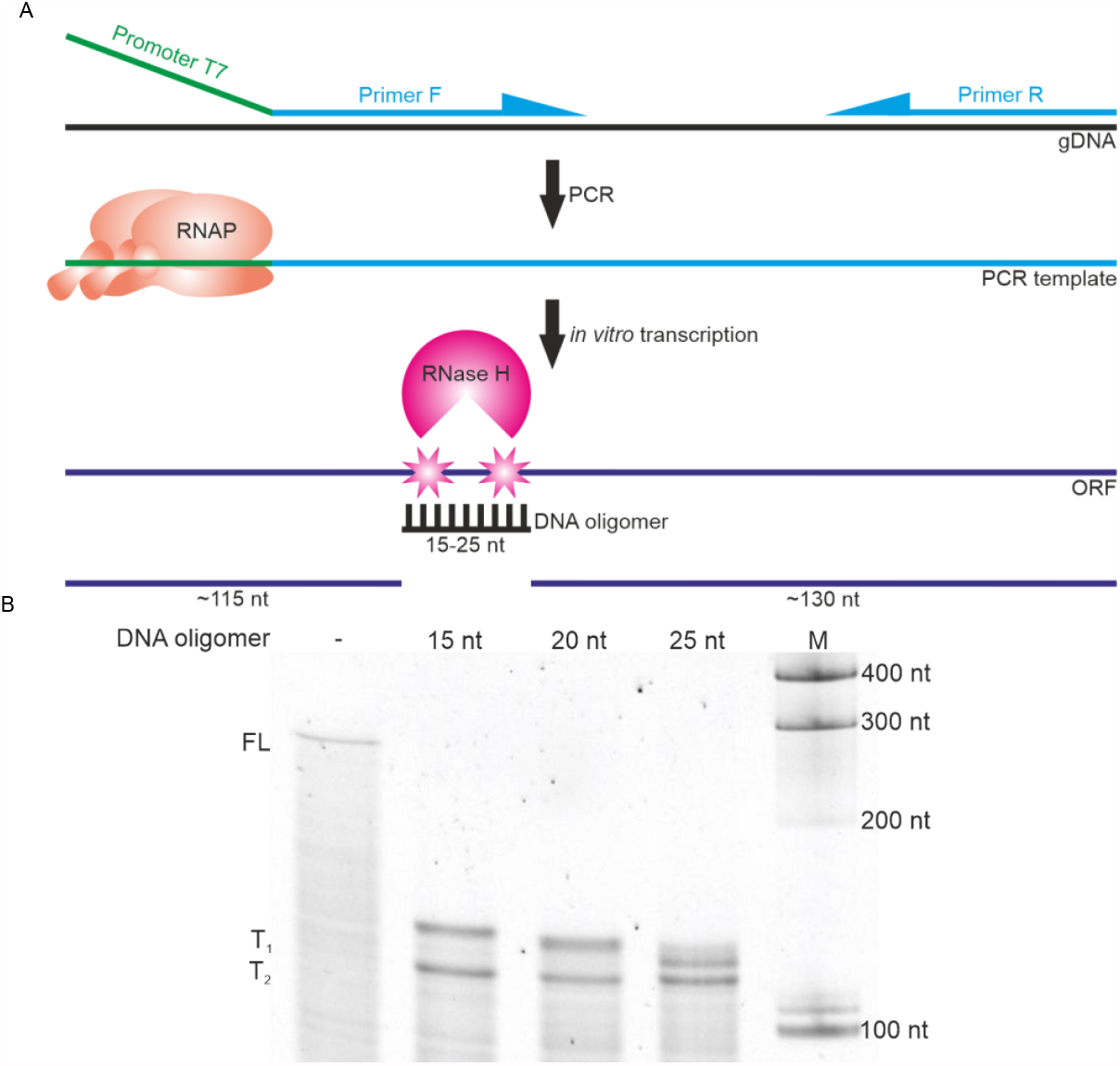
RNase H cleavage of *gyrA in vitro* transcripts with different DNA oligomers: 15 nt, 20 nt and 25 nt long. Schematic representation of RNase H nucleolytic activity for *in vitro* trasncripts (A) and the experimental results of RNase H activity (B) M – RNA size marker, FL – full-length transcript, T_1_ and T_2_ – products of RNase H cleavage. „-” – control sample without DNA oligomer.

In the next step, the concentration of the DNA oligomers was optimized, so that the reaction proceeded in nearly 100% efficiency. The results indicate that most efficient cleavage by RNase H occurs at DNA oligomer concentration of 100 nM (Fig. 4, lane 7). Thus, the optimal DNA:RNA molar ratio was estimated as 1:10. Further increase in DNA oligomer concentration caused a gradual degradation of the proper cleavage product (T_1_) and an appearance of a shorter product (T_2_), probably due to unspecific binding of the oligomer induced by its high concentration.

**Figure 4.**
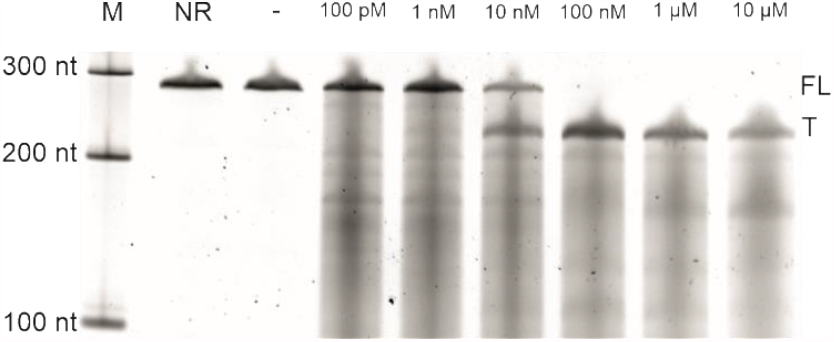
RNase H cleavage of *metE in vitro* transcripts in growing gradient of DNA oligomers. M – RNA size marker, NR - no reaction, “-” – control reaction with no DNA oligomer, FL – full-length transcript, T1 and T2 - RNase H cleavage products.

### *Ex vivo* transcript are efficiently cleaved by RNase H

Previous experiments proved that the designed DNA oligomers are able to efficiently and precisely determine the RNase H site of *in vitro* synthesized transcripts. However, the actual purpose of the proposed method was the analysis of natural transcripts in the pool of total RNA extracted from cells (called here *ex vivo* transcripts). To provide sufficient sensitivity, for visualization we have employed the RT-PCR reaction using three pairs of PCR primers targeted towards 5’, and 3’ parts, as well as the cleavage site (Fig. 2). In the first step, a wide range of DNA oligomer concentrations was tested, from 10 nM to 5 µM (Fig. 5A, 5B). The observable amount of the PCR product amplified by primer set 2 (flanking the cleavage site) were significantly reduced starting from DNA oligomer concentration of 0.1 µM. It suggests the efficient cleavage of a control *gyrA* transcript in the pool of *ex vivo* transcripts by RNase H. Simultaneously, a general lack of changes in intensity of signals derived from products amplified by starters 1 and 3 (up- and downstream cleavage site, respectively), indicated that RNase H cleavage of a *gyrA* transcript in the pool of *ex vivo* transcripts was site-specific. PCR product amplified by starter sets 1 and 3 showed a gradual decrease as the concentration of DNA oligomer increases, starting from a concentration of 500 nM, until a complete disappearance at 5 µM (Fig. 5A). In lanes with the highest DNA oligomer concentrations band signal disappeared, probably due to the high concentration of DNA oligomer, which might interfere with cDNA synthesis or PCR amplification.

**Figure 5.**
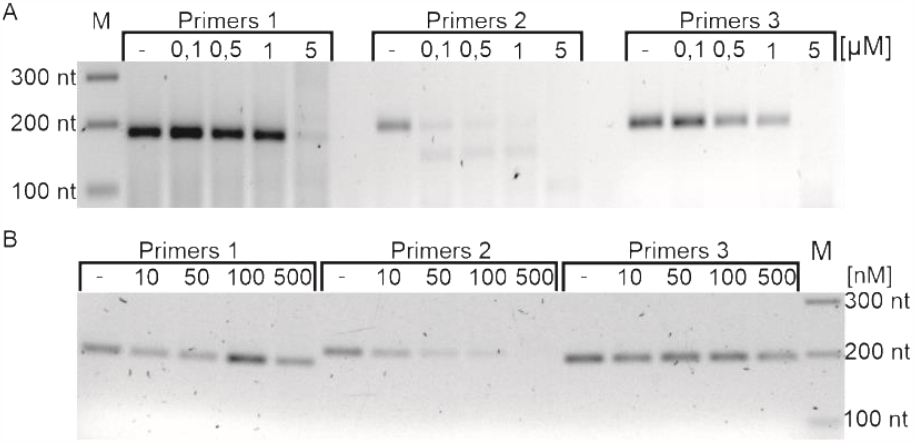
Products of RNase H cleavage of a control *gyrA* transcript in the pool of *ex vivo* transcripts.

Based on above results, for further work we have chosen a range of oligomer concentrations, from 50 to 250 nM, for which changes in cleavage efficiency were observed. In the next step, we have compared the influence of oligomer concentration on cleavage of SAM riboswitch-containing mRNAs: *samT, metIC, metE, mtnKA*. For all tested transcripts, the optimal concentration was in range of 50-100 nM, similar to *gyrA* transcript (Fig. 6). We observed a disappearance of signals from products amplified by primers 2 already after addition of the lowest DNA oligomer concentration of 50 nM. The signals derived from PCR products of primers 1 and 3 were not significantly decreased even with increasing DNA oligomer concentration, except of *mtnKA* and *metE* transcripts, where reduced signal was observed only with 250 nM DNA oligomers.

**Figure 6.**
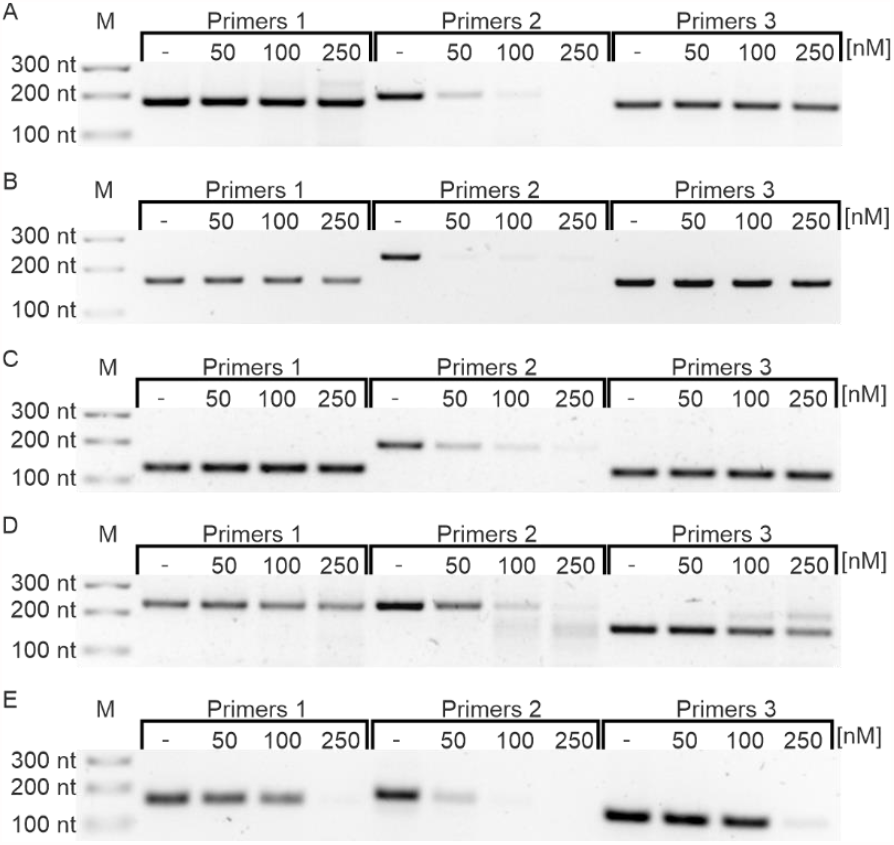
Products of RNase H cleavage of a riboswitch-containing transcripts in the pool of *ex vivo* transcripts. *gyrA* (A), *samT* (B), *metIC* (C), *metE* (D), *mtnKA* (E), M-DNA size marker.

We concluded that the most optimal concentration of DNA oligomer depends slightly on the gene/operon and equals to 50 or 100 nM per 100 ng of total RNA. These are the conditions for which a significant reduction (or a disappearance) of the amplification signal of primers 2 and the permanence of signals from primers 1 and 3 are simultaneously observed.

### ddPCR allows for quantitative analysis of RNase H cleavage of *ex vivo* transcripts

Because the analysis of PCR products has, at best, a semiquantitative character, we introduced a quantitative ddPCR analysis to the proposed protocol..

We performed RNase H cleavage reactions of gyrA, samT, metE, metIC and mtnKA transcripts in the pool of *ex vivo* transcripts followed by ddPCR (Fig. 7). We observed a visible reduction in primers 2 amplification product concentrations in all tested transcripts already after adding the smallest amount of DNA oligomer (50 nM). Interestingly, when applying ddPCR to quantification 5’ and 3’ parts of the cleaved transcripts (primer sets 1 and 3), we have not observed the decrease of products with increase of oligomer concentration to 250 nM, as it was in classical RT-PCR. This observation strongly support the superiority of ddPCR-based quantification over classical RT-PCR methods. Furthermore, the employment of ddPCR allowed for a precise quantification and calculation of RNase cleavage efficiency. The average cleavage efficiency oscillated around 83%, 97% and 97% for *gyrA* control transcript and 94%, 96% and 95% for riboswitch-contained transcripts (for DNA oligomer concentrations of 50 nM, 100 nM and 250 nM, respectively).

**Figure 7.**
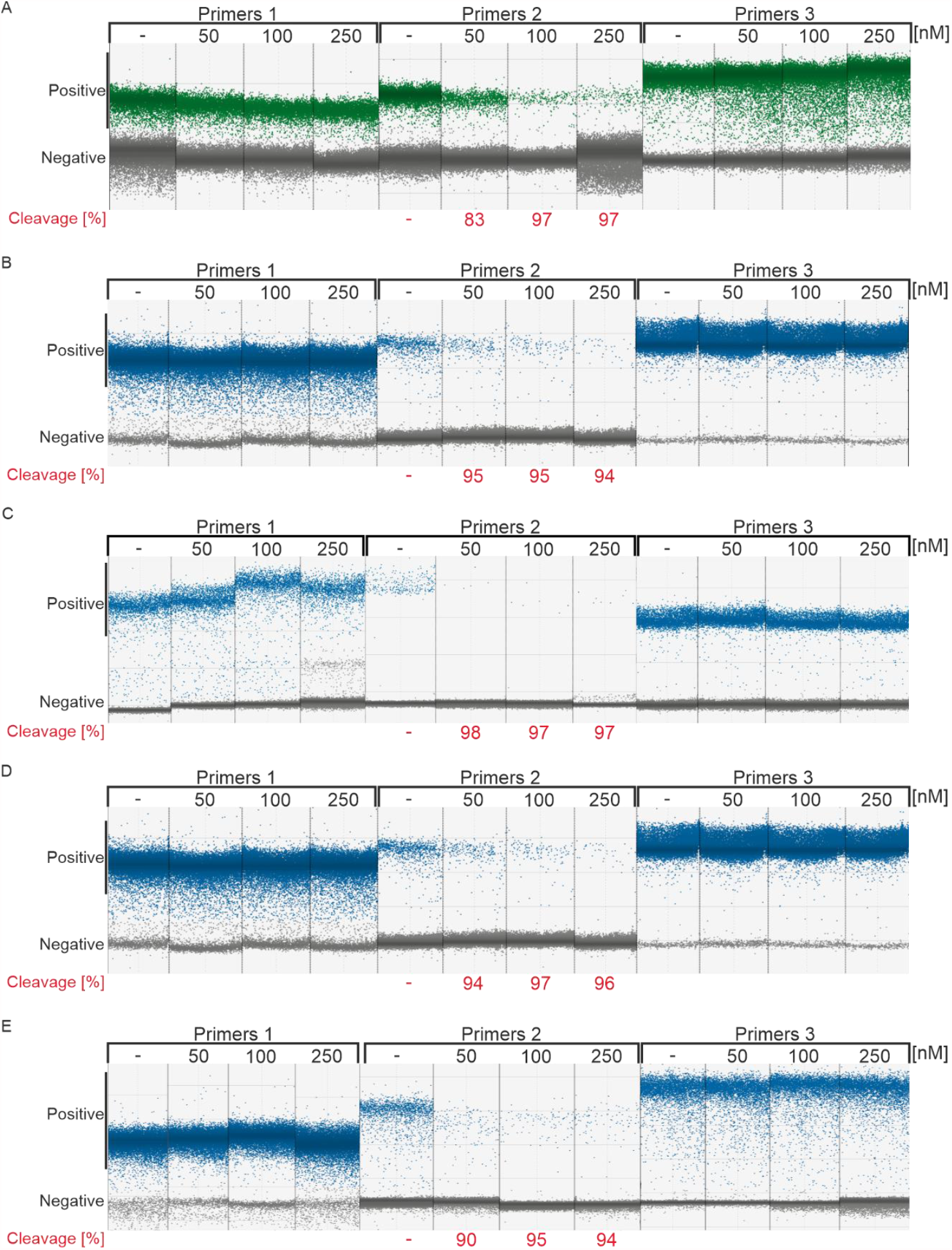
ddPCR result of *ex vivo* transcripts after RNase H cleavage. *gyrA* (A), *samT* (B), *metE* (C), *metIC* (D), *mtnKA* (E).

For each set of primers (primers 1, 2 and 3) DNA oligomer gradient from 0 nM (-) to 250 nM was applied. Positive droplets are marked as green dots, negative - as gray dots. Positive droplets, due to the presence of the EvaGreen fluorescent dye, emit light only with PCR reaction product inside. The higher input concentration of a given transcript, the more copies of the cDNA and the greater the number of positive versus negative droplets.

### RNaseH/ddPCR method allows for determination of SAM riboswitches induction under methionine starvation

Analysis of RNase H cleavage efficiency by ddPCR revealed that 50 or 100 nM of DNA oligomers per 100 ng of total RNA is the optimal amount, as a significant reduction in concentration of amplification products from primers 2, whereas the permanence of the signal from primers 1 and 3 was retained. We have therefore decided to employ our newly established method to study the dynamics of SAM riboswitch activity in *Bacillus subtilis* in a response to progressing methionine starvation. We used 4 different time-points of methionine elimination from bacterial culture, extracted total RNA, performed RNase H cleavage reactions targeted against *ex vivo gyrA* (non-riboswitch control), *samT, metIC, metE* and *mtnKA* transcripts and calculated an absolute quantities of the resulting ddPCR amplification products (Fig. 9 and **Błąd! Nie można odnaleźć źródła odwołania**.).

**Figure 8.**
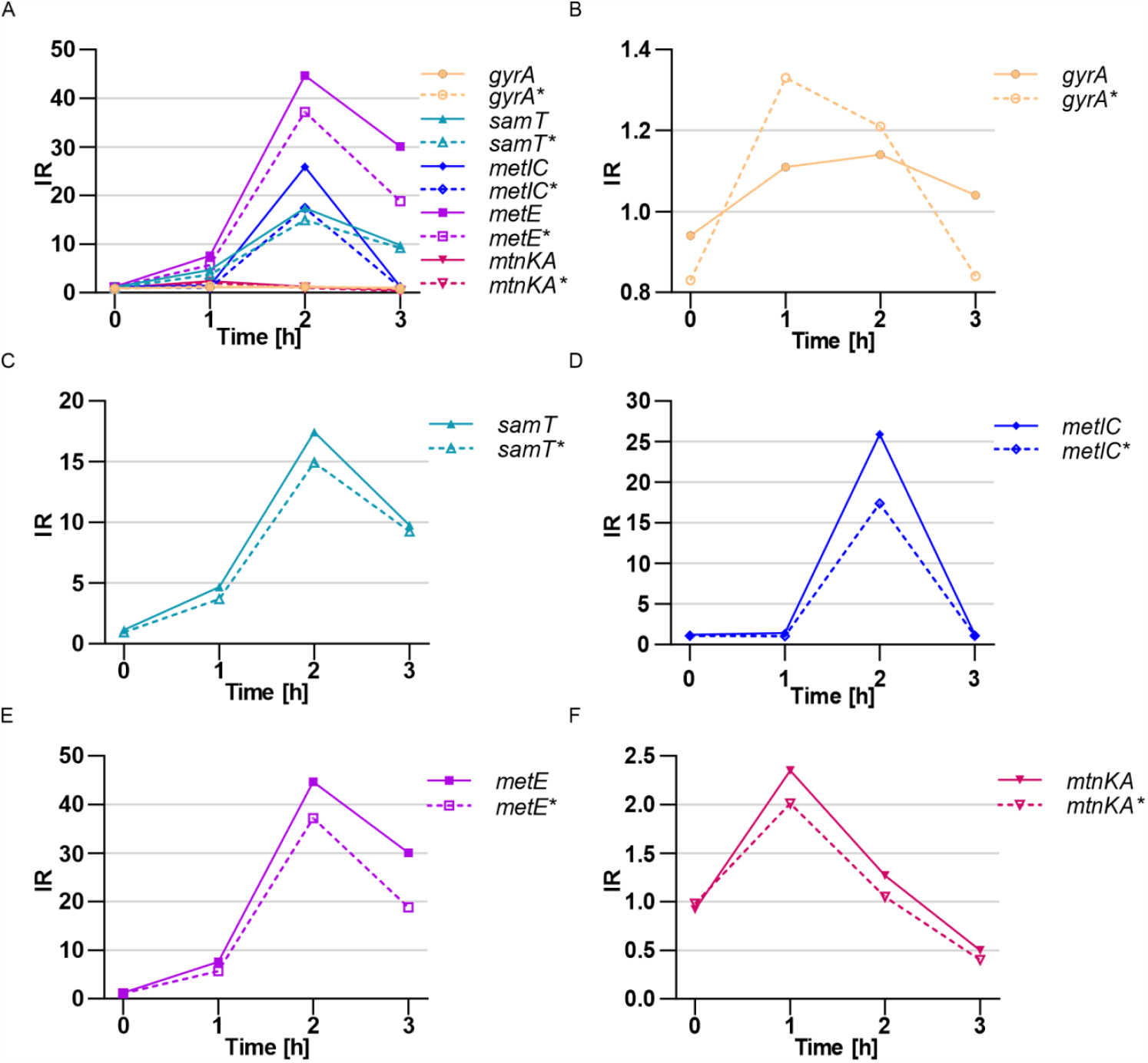
Induction ratio of SAM riboswitches measured by ddPCR. IR – induction ratio. Samples were treated with RNase H cleavage with (solid lines) and without DNA oligomer (dashed lines). Analysis was performed for: combined data (A), *gyrA* (B), *samT* (C), *metIC* (D), *metE* (E) and *mtnKA* (F) in four time points [h].

**Figure 9.**
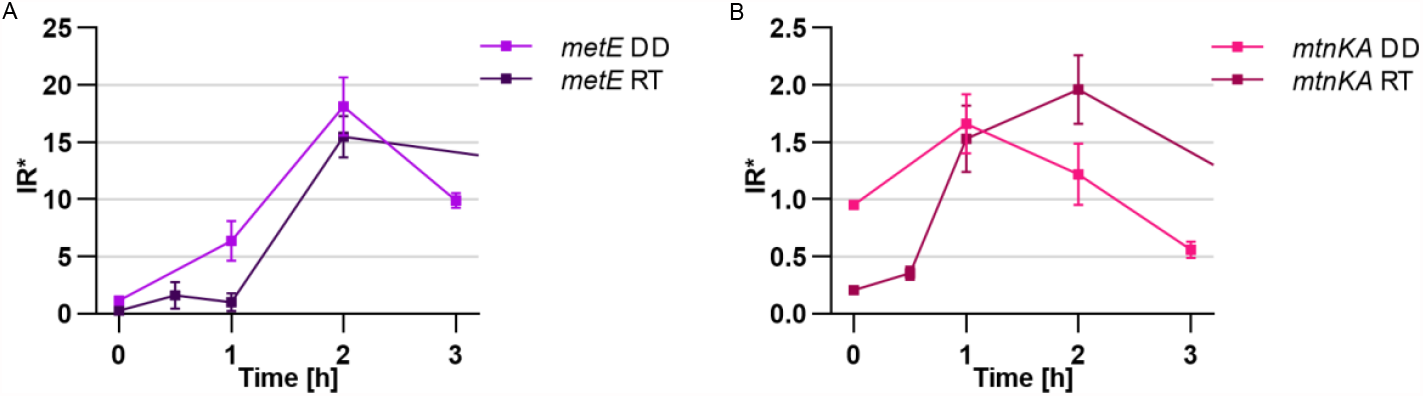
Comparison of IR^*^ between ddPCR and qRT-PCR. ddPCR- brighter colors (DD). qRT-PCR - darker colors (RT). Analysis was performed for two genes: *metE* (A) and *mtnKA* (B).

Analysis of concentration of individual transcripts, expressed in copies/µl, proved that in vast majority of samples the concentration of total number of transcripts (the sum of full length and truncated) was comparable, with minor exceptions, like *metE* gene at 2 h of starvation (**Błąd! Nie można odnaleźć źródła odwołania**.). However, a significant differences were observed in the ratio of full-lenght to truncated transcripts, with a overrepresentation of the truncated transcript in methionine presence and shift towards full-length transcript under methionine starvation, reflecting the activation of the methionine synthesis (Fig.). The largest changes in the quantitative composition are observed at 2 h time point. At 0 h time point (after transfer to a new media), in turn, we generally observed the lowest total expression level, except for *samT* transcript. Thus, for most of the studied genes, the appearance of full-length transcripts seem to be caused by novel transcription under low concentration of methionine which does not activate the riboswitch-controlled PTT (Fig.10). This is particularly visible at 2 h time point for *metIC, metE* and *mtnKA* genes/operons and at 3 h time point for *metIC* and *metE* genes. *mtnKA* operon does not show significant changes in induction ratio, comparable to the control – *gyrA*, suggesting that riboswitch domain detected in its 5’UTR could be acting via translation regulation or being inactive pseudo-riboswitch.

**Figure 10.**
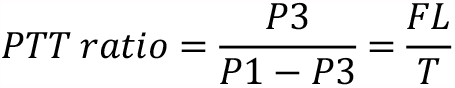
PTT ratio calculation. FL – concentration of full-length transcript. T – concentration of PTT transcripts. P1 – signal from primers 1. P3 – signal from primers 3.

**Figure 10.**
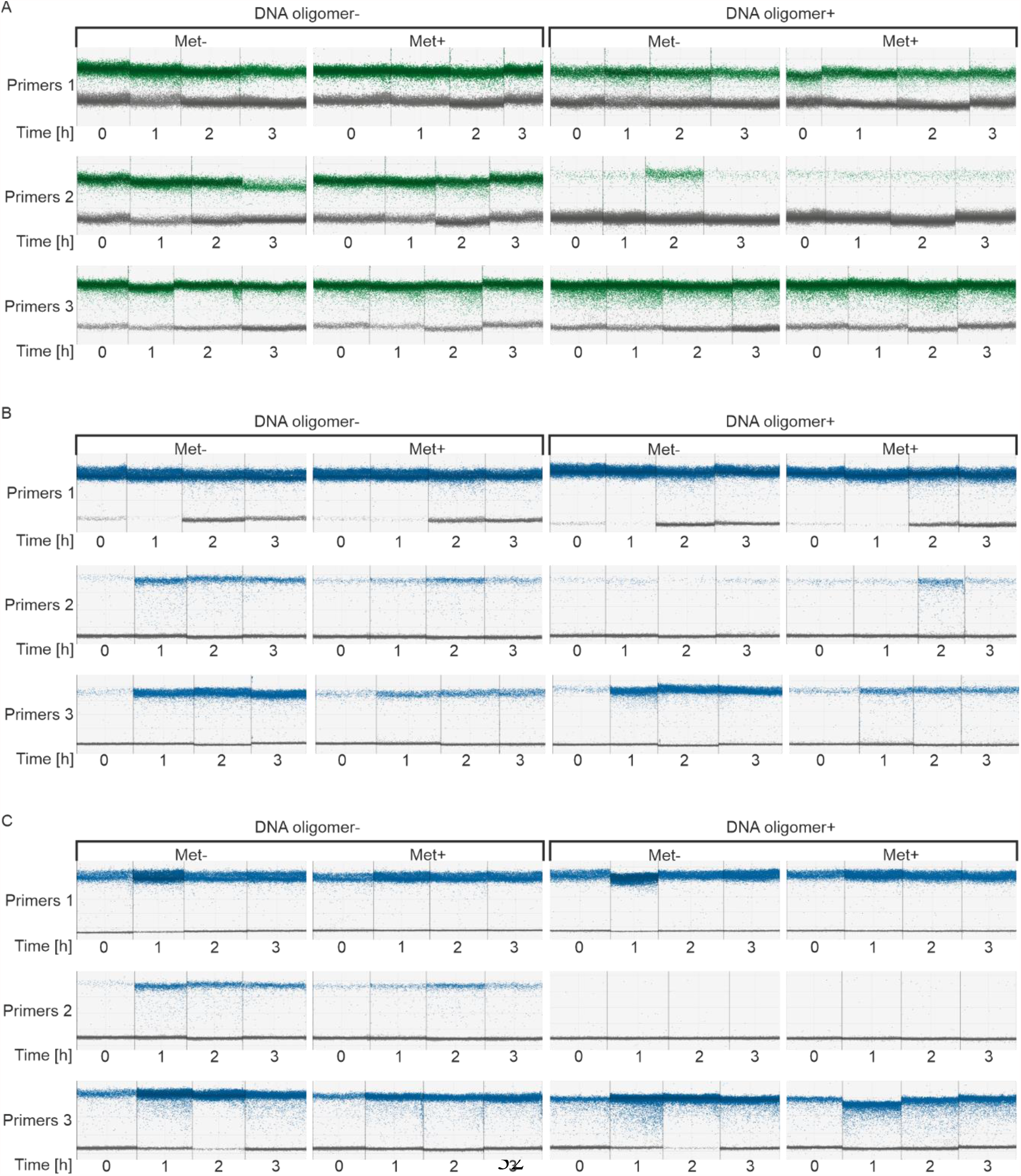
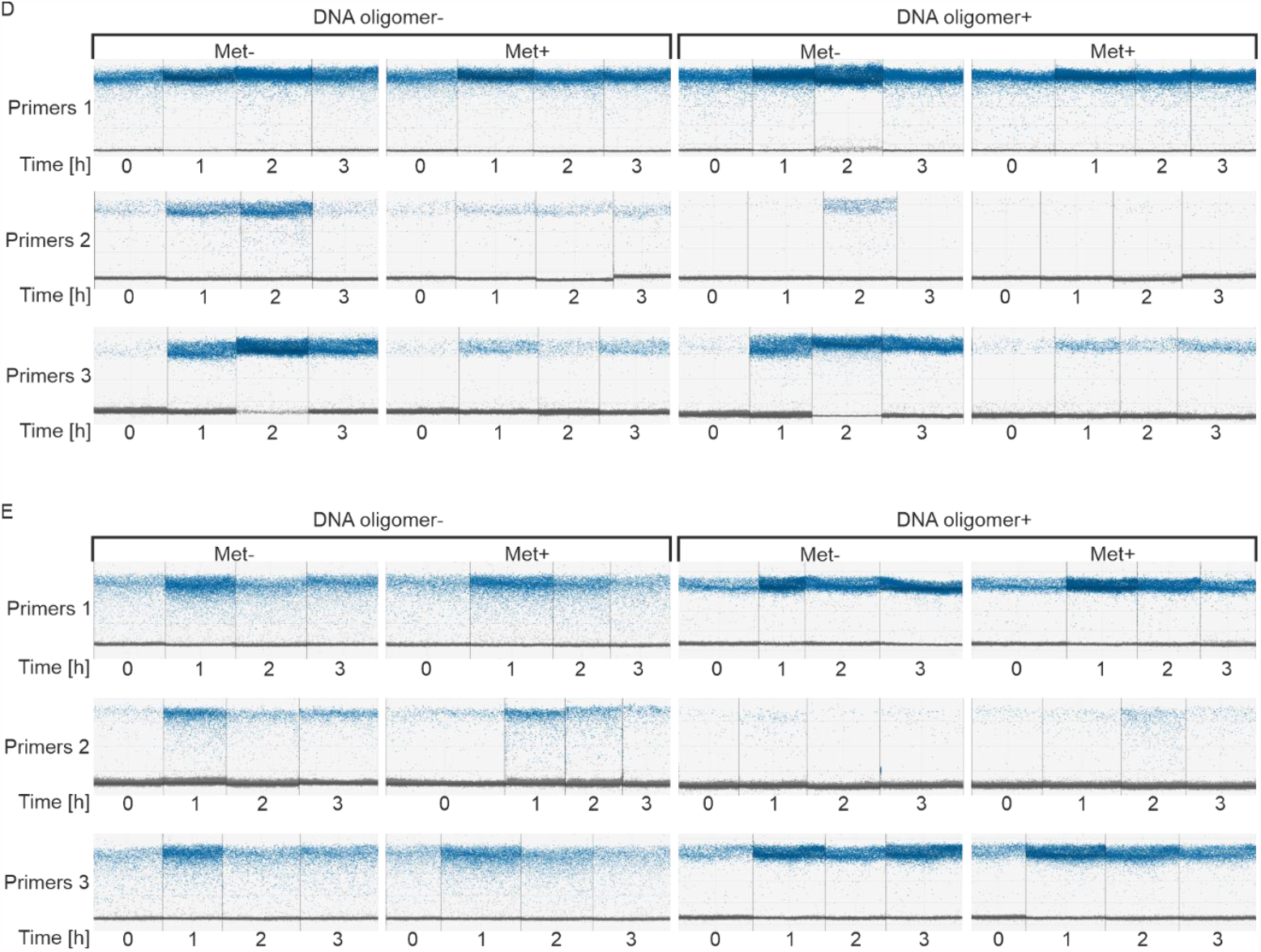
Analysis of transcriptional activity of methionine-induced riboswitches at different time points. (A) control transcript *gyrA*, (B) *samT*, (C) *metIC*, (D) *metE*, (E) *mtnKA*. Time [h] – time of bacteria growth in methionine elimination medium. Met– - lack of methionine in medium. Met + - presence of methionine in medium.

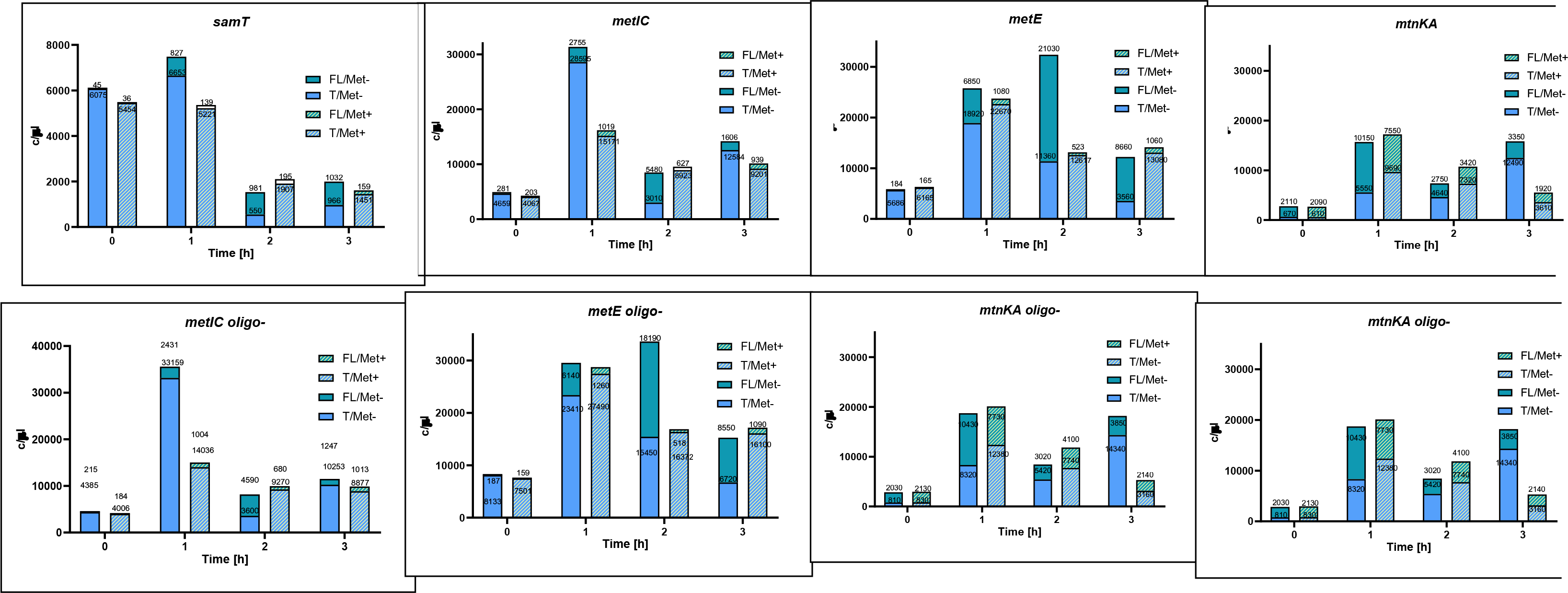

Based on full-length and truncated transcripts concentration from ddPCR analysis, we calculated the induction ratios of SAM riboswitch in *Bacillus subtilis*, defined as a ratio between normalized amount of full-lenght transcripts in *holo* state of the riboswitch (induced by methionine starvation. Met-) to the *apo* state (in the presence of methionine, Met+) (Fig. 10). The induction ratio was estimated separately for four time points.

**Figure 10.**
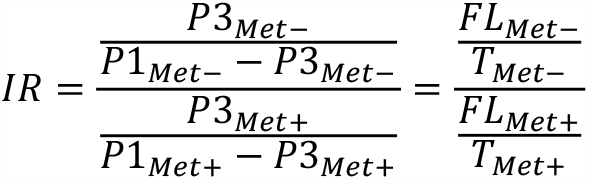
Induction ratio calculation. IR – induction ratio. FL_Met-_ – concentration of full-length transcript in methionine starvation conditions. FL_Met+_ – concentration of full-length transcript in methionine conditions. T_Met-_ – concentration of terminated transcript in methionine starvation conditions. T_Met+_ – concentration of terminated transcript in methionine conditions. P1_Met-_ – signal from primers 1 in methionine starvation conditions. P1_Met+_ – signal from primers 1 in methionine conditions. P3_Met-_ – signal from primers 3 in methionine starvation conditions. P3_Met+_ – signal from primers 3 in methionine conditions.

All analyzed genes/operons which expression is controlled by riboswitches showed a gradual increase in IR, starting from 1 for a 0 h time point. with a peak in 2 h and a final decrease after 3 h (Figure 8A). *mtnKA* operon has a slightly different characteristic, it achieves the highest IR in 1 hour and a gradual decrease afterwards (Figure 8F). The expression of *gyrA* does not show significant induction at any time point (Figure 8B), as expected for a control gene, not regulated by any riboswitch. The highest IR increase was observed for *metE* gene at 2 h time point (IR=45) (Figure 8E), followed by *metIC* operon with IR=26 (Figure 8D) and *samT* with IR=17 (Figure 8C). *mtnKA* operon only to a small extent responded to changes in methionine levels. reaching the highest IR=2.5 after 1 hour of methionine starvation (Figure 8F).

The introduction of the RNase H cleavage to the procedure eliminated the influence of the distance from the 3 end on estimated transcripts concentration (Figure 8 and Table 2). For all genes controlled by riboswitches, the introduction of DNA oligomer and RNase H cleavage resulted in an increase in IR relative to the control, by an average of 23% (depending on the gene and time point, even up to 60%, Table 2).

**Table 1.**
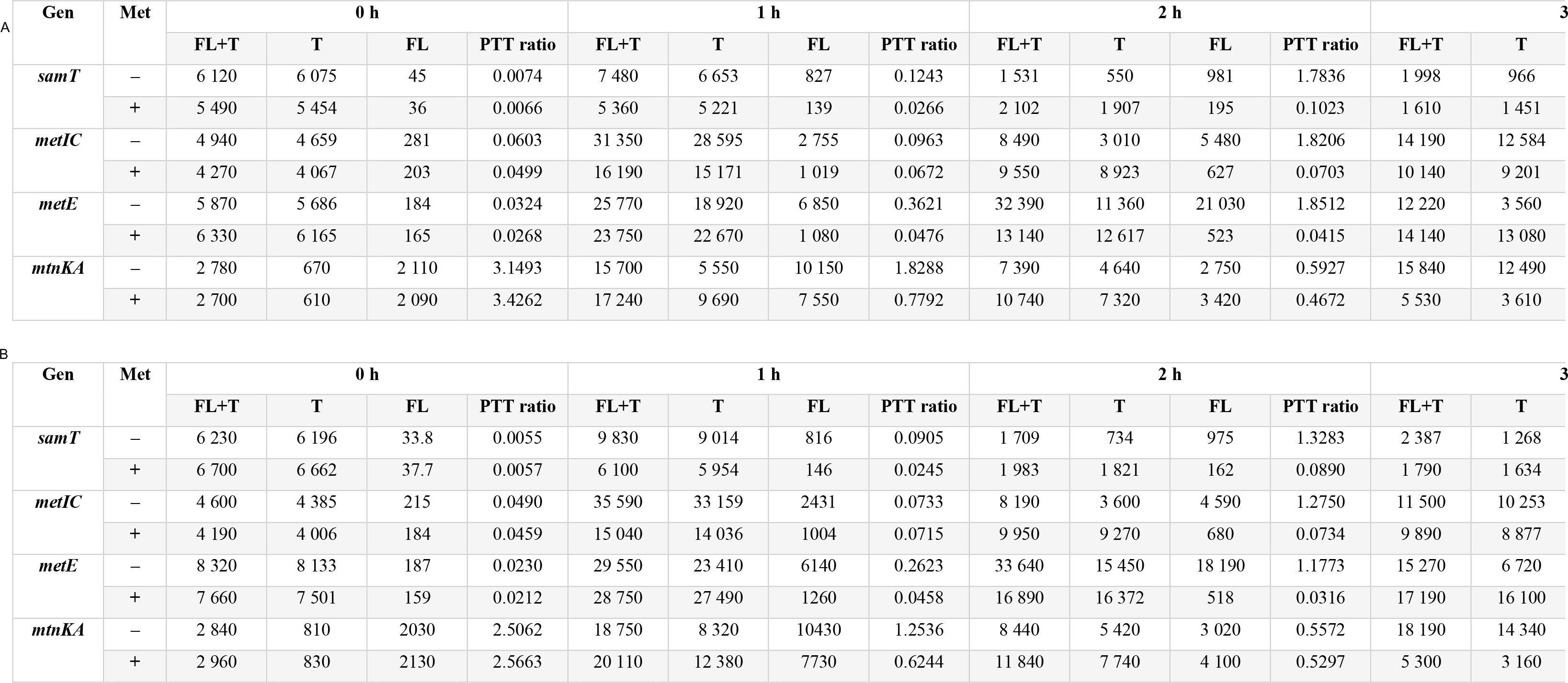
Concentration of riboswitch-containing transcripts at different time points.

**Table 2.** Comparison of induction ratios between samples with and without RNase H cleavage.

| Gene | RNase H cleavage | 0 h | 1 h | 2 h | 3 h |
| --- | --- | --- | --- | --- | --- |
| *gyrA* | - | 0.83 | 1.33 | 1.21 | 0.84 |
|  | + | 0.94 | 1.11 | 1.14 | 1.04 |
| Difference [%] |  | **14** | **-17** | **-6** | **25** |
| *samT* | - | 0.96 | 3.69 | 14.93 | 9.24 |
|  | + | 1.14 | 4.67 | 17.44 | 9.75 |
| Difference [%] |  | **18** | **26** | **17** | **5** |
| *metIC* | - | 1.07 | 1.02 | 17.38 | 1.07 |
|  | + | 1.21 | 1.43 | 25.91 | 1.25 |
| Difference [%] |  | **13** | **40** | **49** | **17** |
| *metE* | - | 1.08 | 5.72 | 37.21 | 18.79 |
|  | + | 1.21 | 7.60 | 44.66 | 30.02 |
| Difference [%] |  | **11** | **33** | **20** | **60** |
| *mtnKA* | - | 0.98 | 2.01 | 1.05 | 0.40 |
|  | + | 0.92 | 2.35 | 1.27 | 0.50 |
| Difference [%] |  | **-6** | **17** | **21** | **27** |

### SAM riboswitch activity with RNaseH/ddPCR is in agreement with qRT-PCR method

Finally, we decided to compare the utility of newly established protocol by comparison of transcriptional activity of riboswitches based on the results obtained using ddPCR method (DD tests) and RT-qPCR method (RT tests). We reasoned that such comparison will allow for the final assessment of the usefulness and adequacy of the newly developed method in relation to the existing one.

We have selected two riboswitches for the comparison: with the highest IR (*metE*) and the lowest IR (*mtnKA*) and performed quantitative real-time PRC analysis (Table 3).

**Table 3.** C_t_ values achieved by RT-qPCR for *metE* and *mtnKA*.

| Gene | Met | 0 h |  | 0.5 h |  | 1 h |  | 2 h |  | 4 h |  |
| --- | --- | --- | --- | --- | --- | --- | --- | --- | --- | --- | --- |
|  |  | FL+T<br>[C<sub>t</sub>]±SD | FL<br>[C<sub>t</sub>]±SD | FL+T<br>[C<sub>t</sub>]±SD | FL<br>[C<sub>t</sub>]±SD | FL+T<br>[C<sub>t</sub>]±SD | FL<br>[C<sub>t</sub>]±SD | FL+T<br>[C<sub>t</sub>]±SD | FL<br>[C<sub>t</sub>]±SD | FL+T<br>[C<sub>t</sub>]±SD | FL<br>[C<sub>t</sub>]±SD |
| *metE* | - | 28.54<br>±0.12 | 33.25<br>±0.48 | 24.51<br>±0.25 | 31.59<br>±0.03 | 23.21<br>±0.23 | 34.43<br>±0.02 | 23.91<br>±0.63 | 31.19<br>±0.17 | 22.03<br>±0.09 | 31.8<br>±0.23 |
|  | + | 29.83<br>±0.16 | 32.63<br>±0.07 | 26.9<br>±0.19 | 34.87<br>±0.54 | 24.8<br>±0.02 | 38.38<br>±0.11 | 26.12<br>±0.09 | 33.43<br>±0.02 | 18.86<br>±0.12 | 32.39<br>±0.69 |
| *mtnKA* | - | 28.53<br>±0.02 | 33.35<br>±0.14 | 24.27<br>±0.15 | 32.13<br>±0.06 | 20.74<br>±0.04 | 34.17<br>±0.2 | 25.55<br>±0.28 | 32.88<br>±0.11 | 25.87<br>±0.18 | 32.02<br>±0.16 |
|  | + | 30.27<br>±0.1 | 32.78<br>±0.07 | 27.73<br>±0.27 | 34.19<br>±0.32 | 20.95<br>±0.01 | 35.15<br>±0.1 | 25.93<br>±0.22 | 33.88<br>±0.16 | 26.9<br>±0.06 | 32.84<br>±0.04 |

Based on the obtained data, we calculated the riboswitch induction ratios (Table 4 and Table 5). However, the induction ratio in RT-qPCR method had to be calculated in a different way than in RNaseH/ddPCR method (Fig. 10), as FL/TT ratio cannot be calculated from RT-qPCR results (only FL/(FL+TT) ratio). The reason is the fact that PTT products cannot be directly quantified with the use of RT-qPCR. So as to compare both methods, new induction ratio (IR*) was applied as shown below (Fig. 12).

**Table 4.**
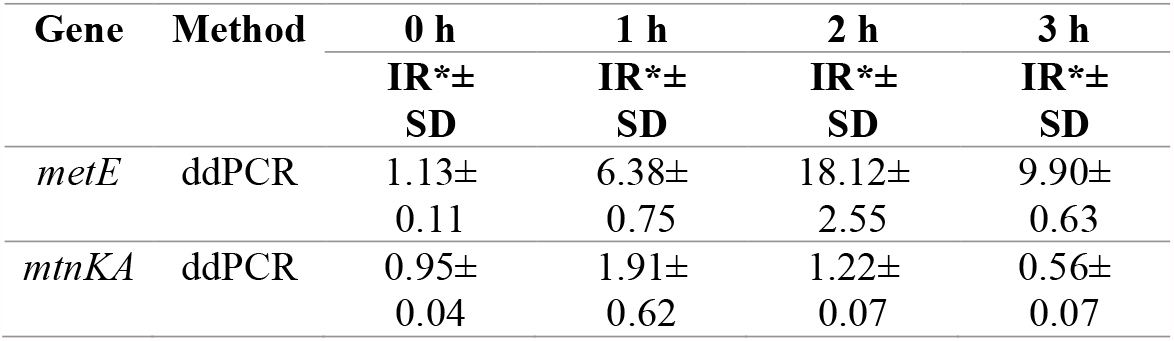
IR^*^ results achieved by ddPCR for *metE* and *mtnKA*.

| Gene | Method | 0 h | 1 h | 2 h | 3 h |
| --- | --- | --- | --- | --- | --- |
|  |  | IR*±<br>SD | IR*±<br>SD | IR*±<br>SD | IR*±<br>SD |
| *metE* | ddPCR | 1.13±<br>0.11 | 6.38±<br>0.75 | 18.12±<br>2.55 | 9.90±<br>0.63 |
| *mtnKA* | ddPCR | 0.95±<br>0.04 | 1.91±<br>0.62 | 1.22±<br>0.07 | 0.56±<br>0.07 |

**Table 5.** IR^*^ results achieved by RT-qPCR for *metE* and *mtnKA*.

| Gene | Method | 0 h | 0.5 h | 1 h | 2 h | 4 h |
| --- | --- | --- | --- | --- | --- | --- |
|  |  | IR*±<br>SD | IR*±<br>SD | IR*±<br>SD | IR*±<br>SD | IR*±<br>SD |
| *metE* | RT-qPCR | 0.27±<br>0.12 | 1.61±<br>1.17 | 1.00±<br>0.79 | 15.50±<br>1.80 | 12.80±<br>4.60 |
| *mtnKA* | RT-qPCR | 0.21±<br>0.04 | 0.36±<br>0.06 | 1.53±<br>0.29 | 1.96±<br>0.30 | 0.87±<br>0.27 |

**Figure 12.**
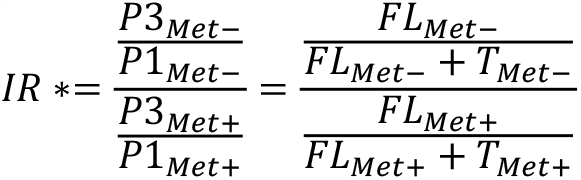
Induction ratio IR^*^ for comparison of ddPCR and RT-qPCR method. IR^*^ – new induction ratio. FL_Met-_ – concentration of full-length transcript in methionine starvation conditions. FL_Met+_ – concentration of full-length transcript in methionine conditions. T_Met-_ – concentration of terminated transcript in methionine starvation conditions. T_Met+_ – concentration of terminated transcript in methionine conditions. P1_Met-_ – signal from primers 1 in methionine starvation conditions. P1_Met+_ – signal from primers 1 in methionine conditions. P3_Met-_ – signal from primers 3 in methionine starvation conditions. P3_Met+_ – signal from primers 3 in methionine conditions.

The dynamic of IR^*^ changes for *metE* gene is very similar for both techniques with increasing IR^*^ up to the highest value of 16 at a time point of 2 h, followed by gradual decrease (Figure 9. A, Tables 4, 5). Particular ddPCR (DD) samples generally reached higher values. In case of *mtnKA* operon, both techniques agreed on the lack of induction of this operon due to the lack of methionine at all time points (Figure 9 B, Tables 4, 5).

### Connection between SAM riboswitch induction profiles and their metabolic function

A thorough analysis of the expression profiles of different genes/operons led us to several characteristic conclusions. Although different genes/operons are controlled by the same class of SAM-I riboswitch. the expression profile over time is different, which refers to both the strength of induction and the time point at which this induction occurs. For example, the expression of *samT* and *metE* is significantly induced in 1 h after methionine depletion (∼4.7 and ∼7.6 fold respectively, Table). Both genes encode methionine syntheses (Fig. 14), directly involved in amino acid production. The early activation of these proteins during methionine starvation might serve as an adaptive mechanism which allows bacteria to quickly compensate for this essential amino acid, protecting against dangerous deficiency. On the other hand, the *metIC* operon is engaged in the biosynthesis of important intermediates in the methionine biosynthesis pathway, cysteine and cystathionine. Due to their indirect role in methionine compensation, the expression is activated at 2 h time point. Among all analyzed genes/operons, the *metE* is characterized by the highest IR in response to methionine concentration (∼45 fold) (Table). It is responsible for biosynthesis of the main and most efficient enzyme producing methionine, which explains the significant increase in its induction. Genes of the *mtnKA* operon respond weakly to the methionine starvation, reaching a peak of IR (∼2.4 fold) (Table) at the 1 h time point. These genes are involved in further steps of the methionine salvage pathway and are not critical to bacterial survival. For that reason their expression does not have to be controlled by SAM concentration as tightly as previous genes/operons.

**Figure 14.**
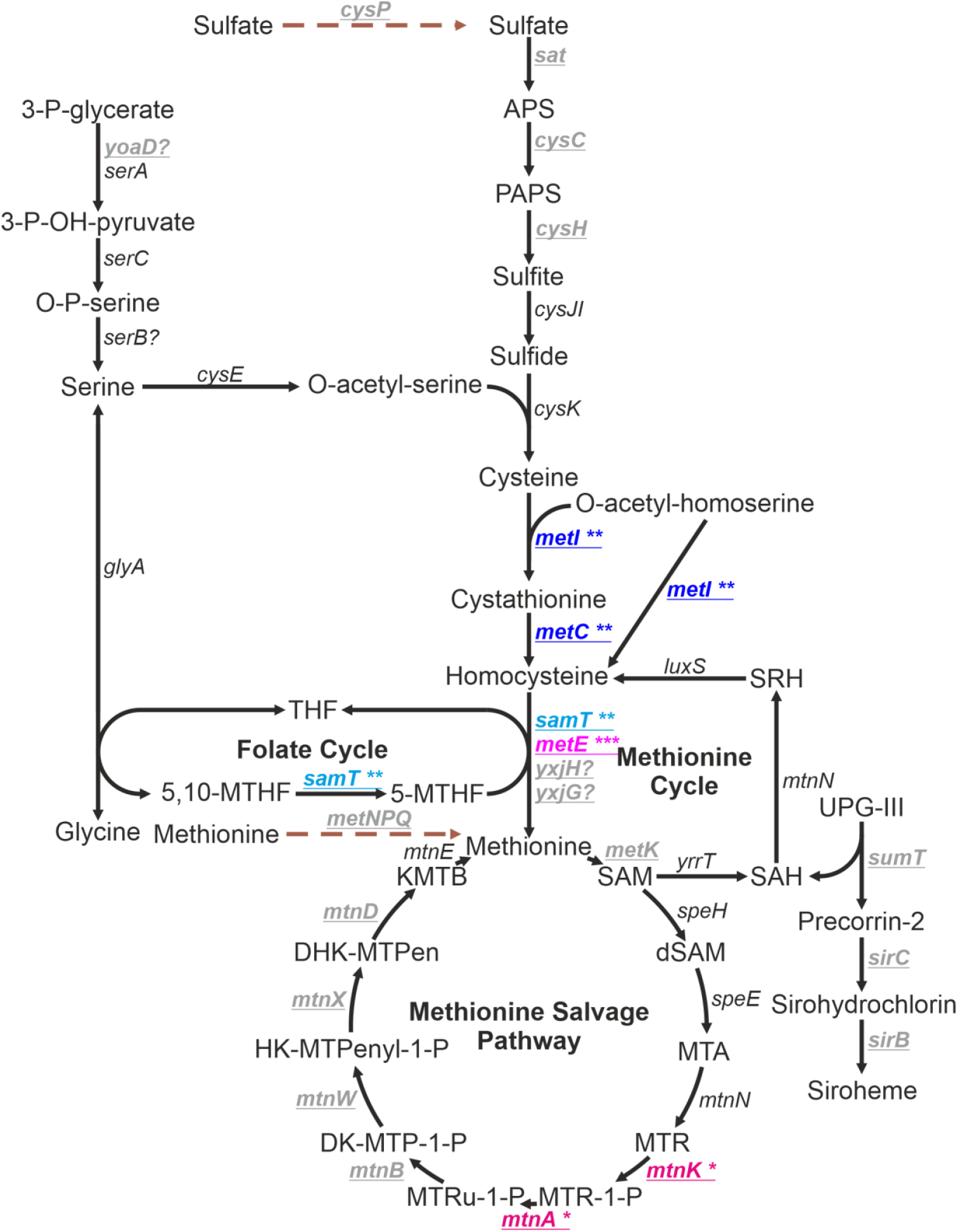
Methionine biosynthesis and metabolism pathway. Underlined and bold and genes are controlled by SAM-I riboswitch. Browned. dashed arrows – methionine import. APS – adenosine-5′-phosphosulfate. PAPS – 3’-phosphoadenosine-5’-phosphosulfate. THF – tetrahydrofolate. MTHF – methylTHF. SAM – S-adenosylmethionine. dSAM – decarboxylated SAM. SAH – S-adenosylhomocysteine. SRH – S-ribulosehomocysteine. MTA – 5′-methyladenosine. MTR – 5′-methylthioribose. MTR-1-P – MTR-1-phosphate. MTRu-1-P – 5-methylthioribulose-1-phosphate. DK-MTP-1-P – 2.3-diketo-5-methylthiopentan-1-phosphate. HK-MTPentyl-1-P – 2-hydroksy-3-keto-5-methylthiopentenyl-1-phosphate. DHK-MTPen-1-P – 1.2-dihydroksy-3-keto-5-methylthiopenten-1-phosphate. KMTB – 2-keto-4-methylthiobutyrate. UPG-III – uroporphyrinogen-III. Asterisks corresponds with induction ratio: ^***^ – high induction, ^**^ – average induction, ^*^ – weak induction.

## Discussion

Taking the advantage of unique properties of the RNase H enzyme and the ddPCR technique, we developed a new method for the quantitative analysis of the transcriptional activity of *Bacillus subtilis* SAM riboswitches. The method proposed herein overcomes the barriers in this process. To date, terminated transcripts could not be easily and precisely quantified; thus, the main attention was focused only on full-length transcripts [23]. Owing to the RT-ddPCR technique, we were able, for the first time, to present absolute concentrations of both full-length and PTT transcripts. The application of RNase H cleavage at the position defined by the DNA oligomer eliminated the impact of the distance from its 3 end on the number of calculated transcripts.

The i*n vitro* and *ex vivo* experiments presented herein proved that RNase H is capable of cleaving any RNA molecule efficiently and specifically, even in a pool of total RNA. It is worth noting that mRNA constitutes only ∼5% of the total RNA of bacterial cells, not to mention the specific transcript of interest [34]. For that reason, achieving an acceptable specificity for the reaction was particularly challenging. The use of the RT-ddPCR gave us the opportunity to more accurately determine the cleavage efficiency, reaching almost 100%.

The real challenge for the developed method is to apply it to solve an independent research problem. In this case, an attempt was made to determine the concentration of full-length and PTT transcripts, as well as calculating an improved IR induction ratio of *Bacillus subtilis* SAM riboswitches. In addition, the IR coefficient was determined both for samples subjected to RNase H cleavage (with DNA oligomer) and for control samples (without DNA oligomer), which allowed to trace the impact of the method on the result obtained. It turned out to be significant and the IR value increased by an average of 23% as a result of RNase H digestion. The average IR increase is consistent with the theoretical assumptions. Crossing transcripts reduces overrepresentation of readings from the end of 5, which leads to a decrease in the number of readings from primers 1 (Fig. 10). In addition, this effect is the stronger the greater the proportion of transcripts are FL molecules, which means that the numerator of the equation increases more (for Met-samples) than the denominator (Met+). Similar results were obtained if compared this method to qRT-PCR. The changes of transcription level in the course of time shown certain regularities, reflecting the physiological functions of individual genes/operons in methionine metabolism.

Considering all the benefits of using the proposed method to perform a transcriptional analysis of riboswitches, a noteworthy idea would be to extend it to all transcriptional riboswitches. However, it is worth remembering that this method does not have to be limited for the analysis of only riboswitches. Applicability of this method can be easily expanded to the analysis of any regulatory elements based on transcription termination mechanisms.

## Acknowledgement

This work was supported by the National Science Centre, Poland [UMO-2016/23/N/NZ1/02446 to P. M.]. The work was also supported by the Polish Ministry of Science and Higher Education, under the KNOW programme.

## Author contributions

PM, KB-Z and MZ designed the study. PM, AW and JZ performed experiments. PM analysed the data and wrote the first draft of the manuscript. All authors contributed to the final version of the manuscript.

Conceptualization: P.M. and K.B-Ż. Protocol design: P.M. Optimization of RNAse H cleavage: P.M. and J.Z. Droplet digital experiments: P.M., A.W.-B. and J.Z. Manuscript writing: P.M. and K.B.-Ż., with the input and revision from all authors.

## Supplementary information

**Table S1.**
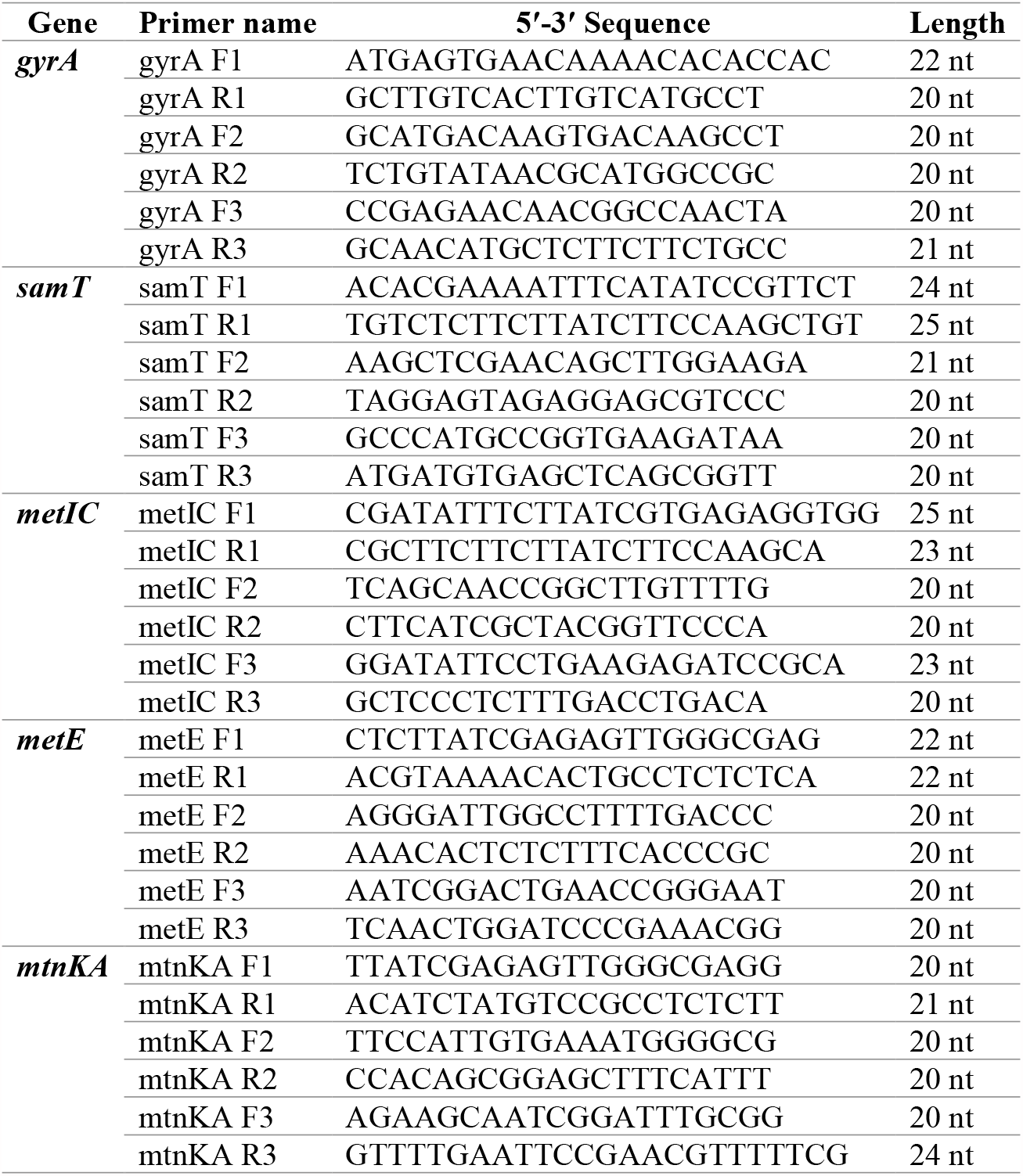
Primers used for PCR, ddPCR and RT-qPCR reaction.

**Table S2.**
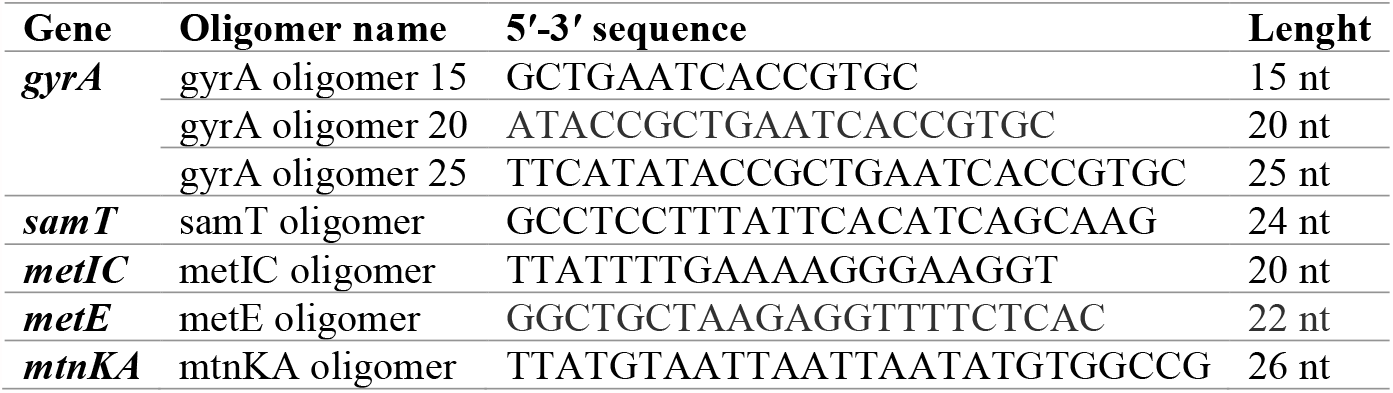
DNA oligomers used for RNase H cleavage reaction.

**Table S3.**
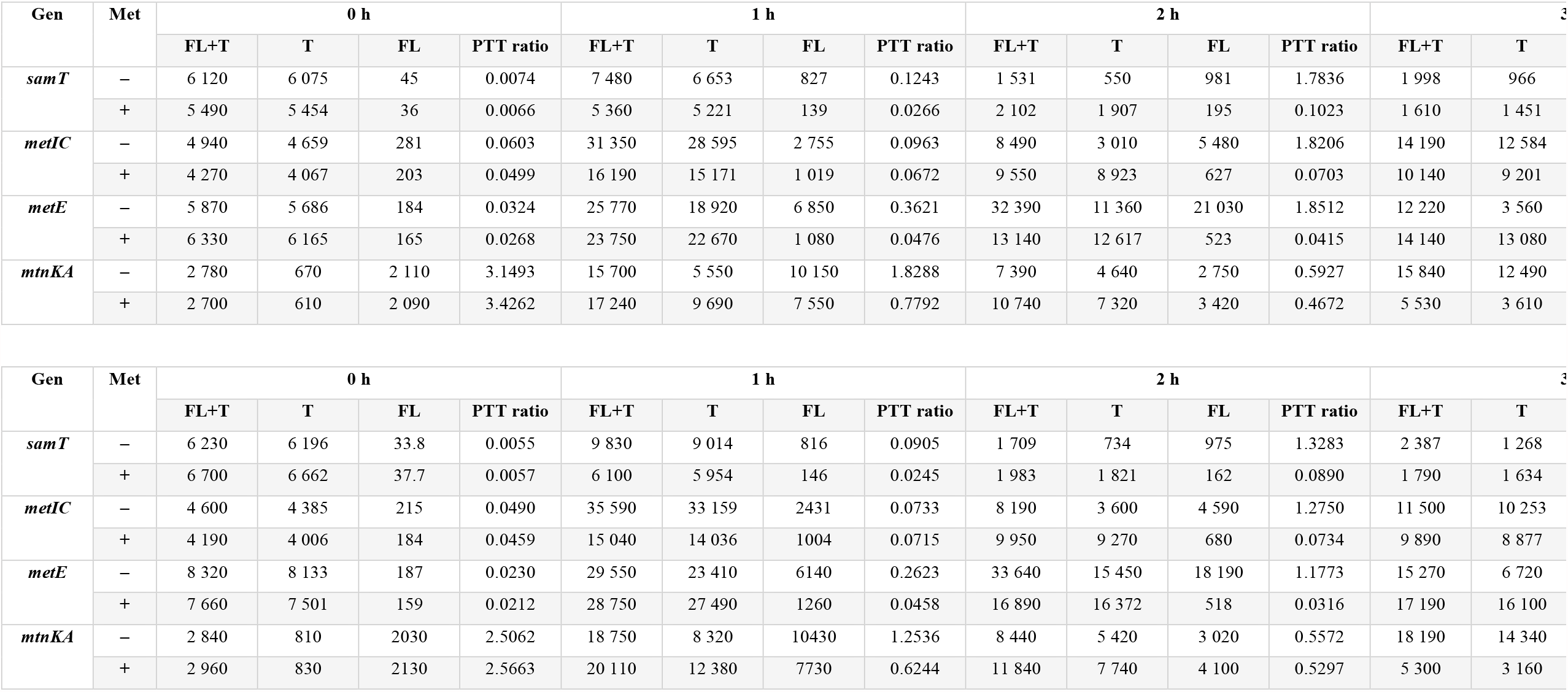
Concentrations and PTT ratio of particular genes/operons achieved by RT-ddPCR in DNA oligomer + (A) and DNA oligomer – (B) conditions.

## Notes

Funding (information that explains whether and by whom the research was supported) This work was supported by the National Science Centre, Poland (UMO-2016/23/N/NZ1/02446 to P.M.).

Conflicts of interest: The authors declare no conflict of interest.

### Competing Interest Statement

The authors have declared no competing interest.

